# Post-acute administration of the GABA_A_ α5 antagonist S44819 promotes recovery of peripheral limb fine motor skills after permanent distal middle cerebral artery occlusion in rats

**DOI:** 10.1101/721282

**Authors:** Marta Pace, Matteo Falappa, Patricia Machado, Laura Facchin, Dirk M. Hermann, Claudio L. Bassetti

## Abstract

**Background:** Ischemic stroke induces hypoexcitability of the peri-infarct cortex by increased tonic GABA activity, which impairs stroke recovery. The GABA_A_ α5 antagonist S44819 has recently been shown to promote post-ischemic motor-coordination recovery after transient proximal middle cerebral artery occlusion (MCAO) in mice. The effects of S44819 on post-ischemic skilled limb movement recovery have so far not yet been assessed in rats.

**Methods:** Male Sprague-Dawley rats were subjected to permanent distal MCAO. Starting 3 days post-stroke, vehicle or S44819 (3 or 10 mg/kg) were delivered orally twice a day for 28 days. A single pellet reaching test was performed at baseline, before treatment onset and at weekly intervals thereafter. Animals were sacrificed at 45 days post-stroke (that is, at 42 days post-treatment onset) after 14 days of drug washout. Body weight was monitored, and infarct size was determined by histology.

**Results:** S44819, administered at 10 mg/kg but not 3 mg/kg significantly improved single pellet reaching performance over 45 days (F(2,96)=22.43, p<0.001). Body weight was not altered. S44819 had no effect on infarct size.

**Conclusion:** Our data indicate that post-acute administration of S44819 at 10 mg/kg promotes skilled forelimb movements. The effect was maintained after S44819 wash out.

## 1. Introduction

GABA (γ-Amino-butyric acid), the predominant inhibitory neurotransmitter in the mammalian central nervous system, has a key role in post-stroke functional recovery. Ischemic stroke causes a hypo-excitability in the peri-infarct motor neo-cortex that results from increased tonic GABA activity onto neurons [1, 2]. Whereas this prolonged elevated inhibition in the peri-infarct region could be useful to protect tissues in the excito-toxic acute phase of stroke, it may impair the neuronal plasticity (re-mapping of peri-infarct regions, neurogenesis, angiogenesis, neurite outgrowth) necessary in a later phase for functional recovery. Therefore, blocking the GABA_A_ receptor responsible for tonic inhibition should promote neuroplasticity and thus improve functional recovery [3, 4].

S44819 is a competitive selective antagonist that interacts with GABA_A_ receptors at the GABA binding site [5]. *Ex vivo* and *in vitro* experiments indicate that S44819 shows preferential binding affinity to α5-containing GABA_A_ receptors [5]. S44819 enhances synaptic plasticity *in vitro* and *in vivo*, improves memory and reduces anxiety in a variety of cognitive tests in rodents [5]. It has recently been shown that post-acute administration p.o. of S44819 at a dose of 10 mg/kg BID enhances neurological recovery and peri-infarct brain remodeling in the post-acute stroke phase in mice exposed to transient proximal middle cerebral artery occlusion (MCAO) [6]. In this study, a set of behavioral tests evaluating motor-coordination and cognitive performance have been studied, but skilled forelimb movements were not assessed. The recovery of fine movements is critical for stroke outcome in human patients.

Following promising phase I studies, which besides others included transcranial magnetic stimulation experiments in healthy humans, in which S44819 increased cortical excitability [7], S44819 is currently examined in a randomized controlled phase IIb trial (Efficacy and Safety Trial with S44819 After Recent Ischemic Cerebral Event [RESTORE Brain]), which investigates the safety and efficacy of S44819 in human stroke patients (https://clinicaltrials.gov/ct2/show/NCT02877615). Considering the importance of fine movements for clinical stroke outcome, we now exposed rats to permanent distal MCAO that induces predominantly cortical brain infarcts [8, 9]. In these rats, we administered S44819 at two doses (3 or 10 mg/kg BID, p.o. starting 3 days post-MCAO) and assessed the effect of S44819 on skilled forelimb movements by the single pellet reaching test.

## 2. MATERIAL AND METHODS

### 2.1 Animals

All experimental procedures were approved by the Animal Research Committee and Veterinary Office of the Canton of Bern (Switzerland). Male Sprague-Dawley rats (8–11 weeks, 300±50 g) housed under 12-hour light/dark cycles (light on 08:00–20:00) at an ambient temperature of 22.0±0.5°C were kept in individual cages. Food and water were provided ad libitum, except for 12-hour intervals prior to behavioral tests, in which animals were food-deprived. All efforts were made to minimize animal suffering and the number of animals used according to the principles of the 3Rs [10]. After a pre-analysis of a set of 5 animals per group, which exhibited statistically significant differences in pellet reaching between groups, further animals were not included in this study.

### 2.2 Experimental design and drug administration

Rats were randomly assigned to three experimental groups who was subjected to MCAO and then treated from 3 to 31 days post-stroke with vehicle (99.5% aqoat, 0.5% magnesium, suspended in 2% hydroxyethyl cellulose) or S44819 (3 or 10 mg/kg; in 30% aqoat milled extrudate, 69.5% aqoat, 0.5% magnesium, suspended in 2% hydroxyethyl cellulose) at two different dosages (3 or 10 mg/kg body weight) (Fig. 1A). Treatments were administered twice daily at 8 AM and 8 PM by oral gavage. After 2 weeks of washout period (day 45) animals were sacrificed. The time window when rats were sacrificed was from 9:00 AM to 12:00 AM after the last single pellet reaching test. The brains were extracted and immediately frozen using dry ice to assess the infarct volume.

**Figure 1.**
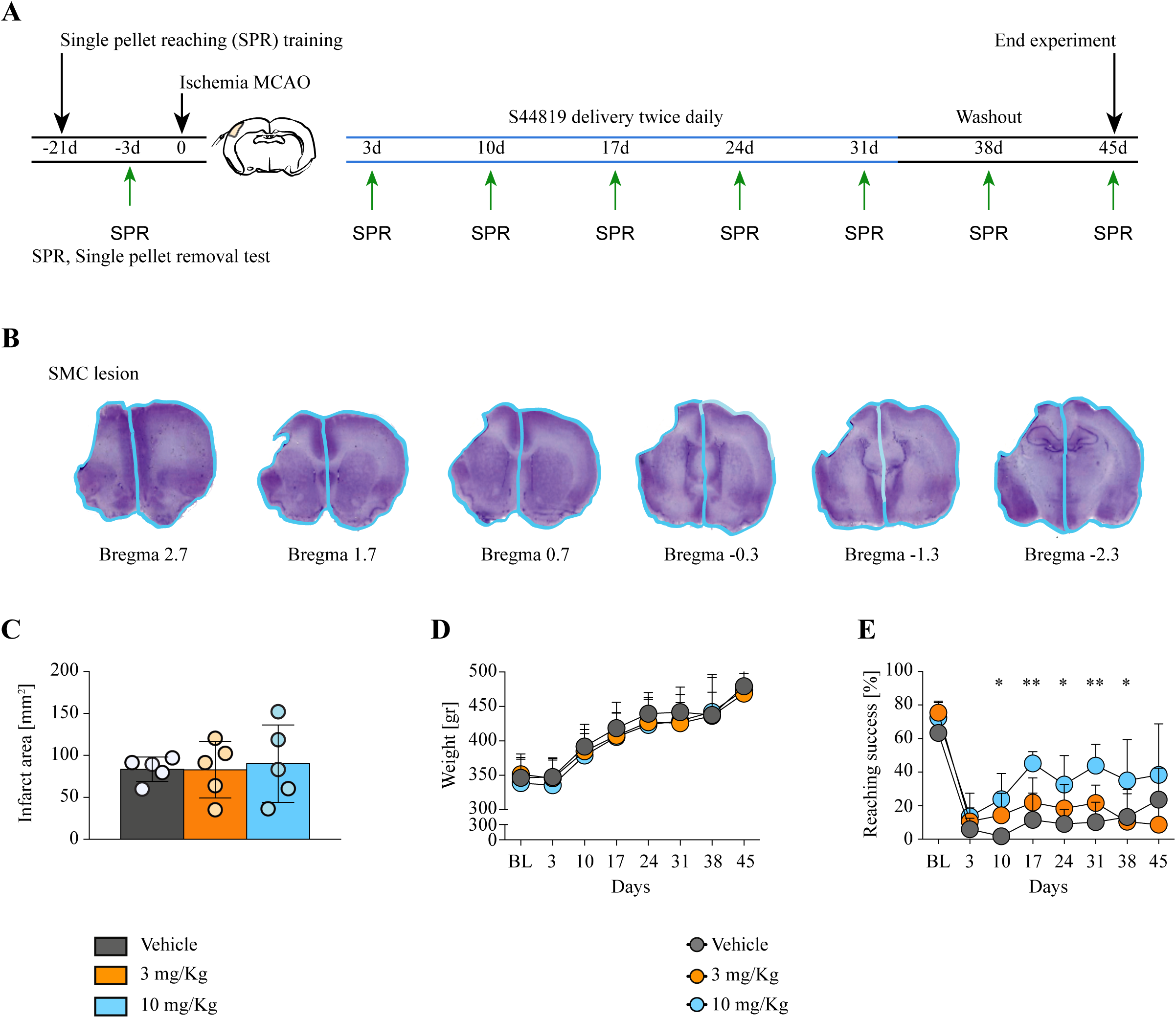
GABA_A_ antagonist S44819 promotes skilled forelimb recovery evaluated by the single pellet reaching test. **(A)** Experimental design: Rats were trained in the single pellet reaching (SPR) test for 3-4 weeks before stroke. From 3 days until 31 days post-MCAO, rats were treated per os with either vehicle or S44819 (3 or 10 mg/kg BID), followed by 14 days S44819 washout until 45 days post-MCAO. Single pellet removal test was assessed over 3 days preceding MCAO (baseline examination) and at 3, 10, 17, 24, 31, 38 and 45 days post-MCAO (that is, at 0, 7, 14, 21, 28, 35 and 42 days post-treatment onset). The green arrows indicate days in which single pellets removal test was performed **(B)** Representative sets of cresyl violet-stained brain sections from a rat subjected to permanent MCAO. The health tissues delineated by a thin cyan-colored line. **(C)** Body weight, **(D)** single pellet reaching performance and **(E)** infarct area in rats exposed to permanent distal MCAO treated with vehicle or S44819 (3 or 10 mg/kg) from 3 to 31 days post-MCAO. Data are shown as means ± S.D. *p<0.05/ **p<0.01 compared with vehicle. Dots represent infarct volume of each animal during each time points.

### 2.3 Single pellet reaching test

The single pellet reaching test was used to assess fine forelimb motor skills. Rats had to use their preferred paw to retrieve a food pellet (45 mg dustless precision pellet; Bio-Serv, Frenchtown, NJ) located on a shelf positioned outside the test chamber. The animals were placed in a clear plexiglas box (41×27×37 cm) with a vertical slit (1×15 cm) in the middle of the front wall, 1 cm above the floor. A 2-cm-wide shelf with small wells was mounted at the front of the slit outside the box wall, on which a pellet was placed on the side contralateral to the preferred paw. To establish a baseline measure of motor performance, rats received daily training sessions for 3-4 weeks on this test. Training sessions comprised the reaching of 50 pellets and the session ended when they made 50 attempts or when 15 min elapsed. Reaches were only considered successful if the rats reached the pellets with the preferred paw, grasped and retrieved the seed, and fed them into their mouth [11]. Performance was defined by: Percent success = (number of successful retrievals/n° of presented pellets (up to 50))*100. The baseline performance was calculated as the average of the 3 days immediately preceding surgery. Animals that failed to reach a baseline level of accuracy of 40% were not exposed to MCAO. Post-MCAO, the test was performed at 3 days prior to S44819 treatment onset and at weekly intervals thereafter (that is, at 10, 17, 24, 31, 38 and 45 days) until animal sacrifice.

### 2.4 Induction of focal cerebral ischemia

Rats were subjected to focal cerebral ischemia induced by the three-vessel occlusion method (3Vo) [9, 12], which predominantly affects the somatosensory cortex. 3Vo combines the permanent occlusion of the distal middle cerebral artery (MCA) by electrocoagulation with the permanent occlusion of the ipsilateral common carotid artery by electrocoagulation and the transient occlusion of the contralateral common carotid artery over 60 minutes with an aneurysm clip. For stroke induction, rats were anesthetized with 2% isoflurane (in 30% O_2_). A small piece of skull overlying the MCA was removed and the dura mater was retracted. The MCA and its three main branches were occluded. Stroke was induced in the hemisphere contralateral to the preferred forelimb assessed by the single pellet reaching test. Body temperature was maintained between 36.5±0.5°C by a heating pad during the surgery.

### 2.5 Evaluation of body weight and infarct volume

Body weight was measured at 3, 10, 17, 24, 31, 38 and 45 days post-MCAO. At day 45 after ischemic surgery, rats were decapitated in deep isoflurane anesthesia. Brains were dissected and frozen immediately on dry ice. Coronal 20 μm sections were cut on a cryostat at six predefined levels (L) with 1 mm interval (L-1: 2.7mm; L-2: 1.7mm; L-3: 0.7mm; L-4: −0.3mm; L-5: −1.3mm and L-6: −2.3mm from bregma) [13, 14] (Fig. 1B). One section from each level was stained with cresyl violet and digitized. To determine infarct area, the healthy tissue of both hemispheres was outlined and tissue areas determined in the lesioned hemisphere subtracted from those in the contralesional hemisphere, as previously described [15].

### 2.6 Statistical analyses

Data were presented as mean ± standard deviation (SD). Effects of S44819 on infarct size were evaluated by one-way ANOVA. Effects of S44819 on changes in body weight and motor performance in the single pellet reaching task were evaluated by two-way ANOVA with repeated measures on both factors (groups x days). Whenever statistical significance was achieved, post hoc Tukey tests were performed. GraphPad Prism6 (GraphPad Prism Software Inc., San Diego, CA) was used for statistical analysis. P values were set at 0.05 to indicate statistical significance.

## 3. Results

S44819, administered at 3 or 10 mg/kg, did not influence body weight (Fig. 1D), but enhanced the recovery of fine motor skills evaluated by the single pellet reaching test (effect of treatment group effect: F(2,96)=22.43, p< 0.001; Fig. 1E). Significant improvements of pellet reaching were noted already one week after S44819 treatment onset (i.e., at 10 days post-MCAO). Improvements persisted after the discontinuation of S44819 treatment. Representative videos of vehicle-treated and S44819-treated rats are shown in the Supplementary Materials section. Infarct area did not differ between groups (Fig. 1C).

## 4. Discussion

By exposing rats to permanent distal MCAO, we herein show that post-acute delivery of the GABA_A_ antagonist S44819, which reverses peri-infarct tonic inhibition [5], promotes the recovery of skilled forelimb movements evaluated by the single pellet reaching test, when administered at 10 mg/kg, but not 3 mg/kg starting at 3 days post-MCAO for 28 days. Our results extend data from a recent mouse study, in which S44819 was administered at the same dosage during the same time-window in mice exposed to transient proximal MCAO (Wang et al., 2018). In this study, improvement of motor-coordination recovery and cognitive performance were evaluated by Rotarod, tight rope and Barnes maze tests. Unlike our here-presented study, the recovery of fine movements was not evaluated. In transient proximal MCAO, ischemic injury predominantly affects the striatum and overlying parietal cortex, whereas specifically the motor cortex is spared. As such, the motor deficits are primarily related to subcortical brain injury, which affects the basal ganglia and descending pyramidal tract axons, as well as to injury to sensory and association cortices. Since the recovery of fine movements is critical for clinical outcome in stroke patients, we now complemented this earlier study (Wang et al., 2018) with the present study in a permanent distal MCAO model in rats. The combined evidence of both studies suggests that the antagonism of GABAergic tonic inhibition promotes motor recovery both under conditions directly affecting motor cortex and conditions affecting sensory and association cortices.

Following transient proximal MCAO, the promotion of motor-coordination recovery by S44819 went along with enhanced structural brain remodeling, that is, reduced secondary neurodegeneration, reduced astrogliosis and reduced brain atrophy after 45 days survival (Wang et al., 2018). By means of infarct planimetry, we have been unable in the present study to detect structural tissue preservation effects that translated into reduced infarct area. Differences in the ischemia model (permanent vs. transient MCAO) may explain the different observations between this previous and the present study. Unlike the earlier study, we did not perform in depth histochemical analyses of brain parenchymal remodeling. The present study may also have been underpowered to detect modest effects of S44819 on infarct size. Yet, the expected mode of action of S44819, the promotion of peri-infarct neuroplasticity, is independent of structural survival-promoting effects. The main goal of this study was the analysis of skilled forelimb movements, which can be assessed more reliably in rats than mice. With the detection of recovery-promoting effects in permanent distal MCAO in rats and transient proximal MCAO in mice, the two studies fulfill two key requirements of pre-clinical research, that is, the evaluation of drug effects in at least two species and two stroke models [16]. Although the results of the current study replicate and strengthen data from the previous mouse study [6], an important limitation of this study is its small simple size. Highly significant group differences were noted in this study supporting the efficacy of S44819 in promoting neurological recovery in the postacute stroke phase. The results of the RESTORE-Brain trial, which are expected during summer 2019, are eagerly awaited.

## Author contributions

MP, CB, PM and DMH designed the study. MP performed the animal experiments, assisted by LF. CB provided infrastructural support. MP and MF analyzed the data. MP, DMH and CB drafted and finalized the manuscript. All authors revised and finalized it.

